# Multi-Generation Ecosystem Selection Of Rhizosphere Microbial Communities Associated With Plant Genotype and Biomass In *Arabidopsis thaliana*

**DOI:** 10.1101/2023.09.06.556126

**Authors:** Nachiket Shankar, Prateek Shetty, Tatiana C Melo, Rick Kesseli

## Abstract

The role of the microbiome in shaping the host phenotype has emerged as a critical area of investigation, with implications in ecology, evolution, and host health. The complex and dynamic interactions involving plants and their diverse rhizosphere microbial communities are influenced by a multitude of factors, including but not limited to soil type, environment, and plant genotype. Understanding the impact of these factors on microbial community assembly is key to yielding host-specific and robust benefits for plants, yet remains challenging. Here we ran an artificial ecosystem selection experiment, over eight generations, in *Arabidopsis thaliana* L*er* and Cvi to select soil microbiomes associated with higher or lower biomass of the host. This resulted in divergent microbial communities, shaped by a complex interplay between random environmental variations, plant genotypes, and biomass selection pressures. In the initial phases of the experiment, the genotype and the biomass selection treatment have modest but significant impacts. Over time, the plant genotype and biomass treatments gain more influence, explaining ∼40% of the variation in the microbial community composition. Furthermore, a genotype-specific association of a plant growth-promoting rhizobacterial taxa, *Labraceae* with L*er* and *Rhizobiaceae* with Cvi, is observed under selection for high biomass.

## Introduction

The conventional understanding of the host phenotype involves genetics and environment shaping observable traits. Yet, the last two decades have underscored the microbiome’s significance in shaping the host phenotype, driven by extensive research on its role in ecology, evolution, and host health. The plant microbiome represents a rich source of functional diversity that is not encoded within the host genome. The interactions between the plant host and its microbiome are dynamic and reciprocal, with the plant shaping its immediate environment by exuding specific metabolites, thereby promoting the growth of specific microbial taxa, while the microbiome in turn influences plant health and growth [1]. A multitude of biotic and abiotic factors influence the dynamic nature of the microbiome, including but not limited to plant root exudates [2–4], soil type [5–9], environment [10], and various aspects of the plant, including species [11], genotype [4,12–15] and developmental stage [16,17].

Previous studies in *Arabidopsis thaliana* have shown that the microbial diversity of the soil reduced with proximity to the endophytic compartment, and microbes found in the bulk soil were not enriched in the endophytic compartment and vice-versa [18–20]. Furthermore, the microbiome of greenhouse-grown plants was similar to that of the field-grown plants, in the different plant compartments – rhizosphere, woody stem, and endophytic compartment, indicating an active role of the plant host in creating and maintaining an environment where certain microbes have improved fitness [21]. The soil environment was a stronger predictor of rhizosphere microbiome structure, though plant genotype had a weak influence in some studies [20,22,23]. A follow-up study confirmed that variation in microbiome communities depends more on the environment than on different *Arabidopsis* varieties and sister species. Though there were some differences between different host species, the core microbiome interacting with the plant host at the root endophytic zone was largely consistent and reproducible [24].

Unfortunately, most prior studies were conducted in a single plant growth cycle, precluding an analysis of the temporal stability of the plant-microbiome association. Observations from plant-soil feedback studies highlight that plant-microbiome interactions can be altered by successive growth cycles of a given plant in the same soil [25,26]. Prior studies have employed plant-mediated selection on the soil ecosystem over multiple generations, and resulted in consistent effects on plant characteristics such as biomass, flowering time, and germination [27,28]. By selecting for soils where hosts show the desired phenotype, such as high biomass or altered flowering time, it is possible to enrich microbes that modulate host traits. Additionally, studies showed the plant’s response to abiotic stress, such as drought and salt stress, could be influenced in two ways. First, through generational selection, where plants with desirable stress responses were chosen over multiple generations [29–31]. Second, by introducing beneficial microbes that were associated with plants and had previously lived in similar drought or salt conditions [32]. Mueller et al., [29] used generational selection to create microbial communities that could improve plant seed production by up to 205%. Soil inoculum obtained from drought-exposed soils improved wheat biomass under drought conditions [32]. Older studies, such as Swenson et al. [28], also have shown similar results under optimal growing conditions without necessarily exploring the microbiome. A recent study conducted on the rhizosphere microbiome of wild and domesticated tomato plants over multiple generations demonstrated an escalating influence of host genotype on the microbiome community [33]. A separate study on the phyllosphere microbiome of various tomato genotypes indicated a declining impact of host genotype across successive generations [34].

Despite several earlier efforts, the development of a robust and host-specific microbial community with long-lasting beneficial effects on the plant host remains a challenge. Using an artificial ecosystem selection experiment, we sought to advance our understanding of how plant genotype, environment, and biomass selection treatment (henceforth referred to as biomass treatment) impact the assembly of host microbial communities. Treatments selecting for the microbial communities of high and low biomass plants did affect the growth of plants in subsequent generations, though unknown environmental factors had substantial effects throughout the experiment. In the first few generations of the experiment, the genotype and biomass treatments played a modest role in shaping the microbial community. Over time, the plant genotype and biomass treatments had increasing influence, explaining ∼40% of the difference in the microbial community composition. Moreover, we observe an enrichment of known plant growth-promoting rhizobacteria (PGPR) in the high biomass treatment, indicating potentially beneficial host-specific interactions.

## Material and Methods

### Multi-generation selection of soil ecosystem

*Arabidopsis thaliana* Cvi and L*er* accessions were cultivated using custom-made “rhizotubes,” (Stuart Morey, Univ. of Massachusetts, Boston, unpublished). The tubes were equipped with a black polyethylene sleeve, which effectively blocked light penetration. This design also facilitated the convenient removal of the plant from the pot, granting full access to the root system. (Supplementary Figure S1).

The potting soil used was PRO-MIX PGX, a commercial mixture comprising 80-90% sphagnum peat moss and small quantities of perlite. It was autoclaved twice for 40 minutes with a 48-hour interval between each sterilization. The sterile potting soil was then sifted through a 3 mm sieve and combined with field soil in a 6:1 ratio for the first generation. The field soil obtained from the Center for Agricultural Research in Waltham, Massachusetts, was composed of 44% sand, 49% silt, and 7% clay representative of agricultural and grassland ecosystems.

The mixture was homogenized using a custom-made cement mixer and attached with sterile bins to avoid cross-contamination. Distilled water was gradually added to achieve a final ratio of 1:2 (water to soil). The soil was incubated at room temperature for two days following inoculation before planting the seeds. All seeds used in the experiment were obtained from a single-parent plant. Before planting, the seeds were treated with a solution of 50% bleach vol/vol and a drop of tween 20 for 10 minutes and rinsed ten times with sterile distilled water. We placed 3-4 seeds in the center of each pot and kept only one seedling per pot after emergence. All plants were grown at 22 °C day/ 18 °C night with a 12/12 hour day-night cycle in a controlled growth chamber. Relative humidity ranged from 35-60% and the light intensity was 96 µE. Fertilizer was not applied to plants.

In the first generation of the experiment, 100 plants of each accession (Cvi and L*er*) were grown separately in individual pots. The plants were arranged in a randomized block design to avoid batch effects. All plants were harvested 35 days after germination. The above-ground portion of the plant, the rhizosphere, and the bulk soil from each rhizotube were separated, ensuring there was no cross-contamination (Supplementary Figure S1). The above-ground part of the plant, stem, and leaves were dried at 70 °C for 4 days. All the plants were weighed individually on a closed weighing scale accurate to 1 mg. The root-soil complex (comprising the rhizosphere and the endosphere, henceforth referred to as the rhizosphere) was obtained by shaking the excess soil off the root and placing the root and remaining attached soil in a sterile 5 ml tube. Bulk soil samples were moved to a sterile Ziplock bag. Tubes and bags were immediately transferred to dry ice and then stored at –80 °C for DNA analysis.

All subsequent generations were inoculated using the previously collected bulk soil from the top five and bottom five plants by above-ground biomass. This resulted in two different treatments for each genotype: high biomass L*er*, low biomass L*er*, high biomass Cvi, and low biomass Cvi. This process is repeated for the remaining seven generations. Starting with the third generation, plants were grown in uninoculated sterile potting soil, as a control for random environmental variation (REV). In generation 6 all plants died for unknown reasons. The experiment was then restarted using soil from generation 5. Soil sourced from the top 6-10 plants and the bottom 6-10 plants based on above-ground biomass was used as inoculum to replant generation 6.

### DNA extraction and 16S rRNA amplicon library prep

Microbial DNA was isolated from the frozen rhizosphere samples using the Machery-Nagel Nucleospin Soil DNA extraction kit (MACHEREY-NAGEL Inc., Allentown, PA, USA). Approximately 0.1g of rhizosphere soil sample was used for DNA isolation. All samples were diluted to 5 ng ul-1 with PCR-grade water. 16S rRNA gene was amplified from the isolated DNA samples in triplicate in 96-well PCR plates. The PCR primers used were for the 16S rRNA V4 region, 515F (5’-GTGYCAGCMGCCGCGGTAA-3’) (ref. 35) and 806R (5’-GGACTACNVGGGTWTCTAAT-3’) (ref. 36) for downstream paired-end Illumina (Illumina, Inc., San Diego, CA, USA) barcoded sequencing [37]. The PCR cycling conditions were as follows: 94 °C for 3 min; 25 cycles of 94 °C for 45 s, 50 °C for 60 s, 72 °C for 90 s, and final elongation at 72 °C for 10 min. The triplicate amplified samples were pooled and then purified and normalized using the SequalPrep™ Normalization Plate Kit (Invitrogen Corporation, Carlsbad, Canada). Finally, multiplexed paired-end sequencing was carried out in the Illumina MiSeq (SY-410-1003) platform using earth microbiome project (EMP) primers [38].

### Sequence Data Analysis & Statistics

The paired-end sequences obtained from the Illumina MiSeq were demultiplexed with QIIME 2 and converted into individual sequence fastq files for each sample [37,39]. The rest of the sequence processing was carried out in R (Version 4.2.2) using the DADA2 package (Version 1.24.0) (ref. 40). The reads were processed in R using the following command in DADA2; ‘filterAndTrim(fnFs, filtFs, fnRs, filtRs, truncLen=c(145,145), minLen=50, maxN=0, maxEE=c(2,2), truncQ=2, rm.phix=TRUE, compress=TRUE, multithread=TRUE).’ *De novo* OTU (Operational Taxonomic Units) picking was performed using DADA2 (37) which resolves amplicon sequence variants (ASVs) to single nucleotide differences. Chimeras were removed. The taxonomy was assigned using a 100% cut-off for species-level identification with the (2022) GTDB 16S rRNA reference database [41]. The phylogenetic tree was constructed using IQTree [42], with 1000 ultrafast bootstraps and Modelfinder to identify the best model. In this study, the best-fit model was SYM+R10 chosen according to BIC. The data comprising the OTU table, phylogenetic tree, taxonomy table, and sample meta-data were then parsed through R studio using the phyloseq package [43]. Here, it was processed to remove: OTUs identified as mitochondrial or chloroplast, which had less than five reads across all samples, and samples that had fewer than 8000 reads. The packages phytools, phyloseq, microbiome, and ggplot2 were used for further data analysis. All sequence data and metadata can be downloaded from PRJNA1010286.

### Diversity Analyses

Alpha diversity was computed using four metrics, Observed, Shannon, Chao1, and Simpson diversity indices. Beta diversity was calculated using weighted and unweighted UniFrac, and subsequently Principle Coordinate Analysis (PCoA) plots were constructed using the first two axes that explain the most variance. Linear mixed effect models using, *lmerTest* package in R were used to determine changes in alpha and beta diversity over time (lmer (DiveristyMetric, Genotype*Generation*Biomass Treatment, (1|Generation)), with generation as a fixed and random effect. Pairwise comparisons were determined with *emmeans* and adjusted p-value obtained with Tukey.

### Adonis

PERMANOVA was carried out for each generation using the adonis2 test. Both weighted and unweighted UniFrac distance matrices were used to account for the abundance, presence/absence, and phylogeny of the microbiome. The following model was used ‘adonis2(distance.matrix∼Biomass Treatment+Genotype, data = meta, permutations=999)’.

### Neutral Model

To determine the impact of neutral processes like drift and dispersal or by deterministic selective forces such as plant genotype and biomass treatments on microbial community assembly, we carried out the analysis as described in Burns et al. [44], which fits the data to the neutral model for Prokaryotes from Sloan et al. [45]. The following design was used, sncm.fit (spp=generation(n)_Biomass Treatment, pool = generation(n), stats=T), where n is the nth generation and ‘Biomass Treatment’ is high or low biomass samples.

### Differential Abundance

There are several challenges in estimating differentially abundant (DA) taxa in microbiome data, these include high variability in abundance, zero-inflated data, and the compositional nature of the data. Nearing et al. [46] compared several methods across 38 datasets and found that different methods often identified different sets of DA taxa. The DA taxa are estimated using ANCOMBC2 [47], a conservative and robust approach. ANCOMBC2 incorporates bias correction, effectively addressing sampling-specific and sequencing biases present in the data. This feature ensures that the analysis is not skewed by any systematic errors introduced during the sampling or sequencing process. It also conducts a sensitivity analysis for the pseudo-count addition to assess the impact of different pseudo-count values on zero counts for each taxon. The analysis was run on L*er* and Cvi samples, from OTU to Family taxonomic levels with (fixed effect = (Biomass Treatment + Genotype + Generation), group = Biomass Treatment).

## Results

### Plant Biomass

The ecosystem selection experiment was carried out for the trait plant biomass on two different accessions of *Arabidopsis thaliana,* L*er* and Cvi for eight generations. The two selected treatments were high and low above-ground biomass. The absolute values for the mean biomass of plants (n=50) from each genotype and treatment changed substantially through the course of eight generations of the experiment. Despite the stochastic fluctuations in biomass caused by random environmental variation (REV), the high biomass selected lines were always the same or greater than those of the low selected biomass for both genotypes (Figures 1a and 1b). Due to the significant drop in biomass between generations 1 and 2, a sterile potting soil uninoculated control was planted in all subsequent generations to serve as a reference for REV. The uninoculated control shows that the drastic drop in biomass after generations 1 and 2, was not a result of the inoculated soil but other subtle but crucial environmental factors that could not be held constant. Above-ground biomass for each generation in terms of deviations from the mean was plotted to give a clearer representation of the difference in phenotype seen in every generation (Figures 2a and 2b). Significant differences between the low and high biomass treatments become apparent from generation 4 onwards. Plants of all treatments in generation 6 died resulting in a sharp dip, which necessitated repeating that generation (described in the Materials and Methods).

**Figure 1.**
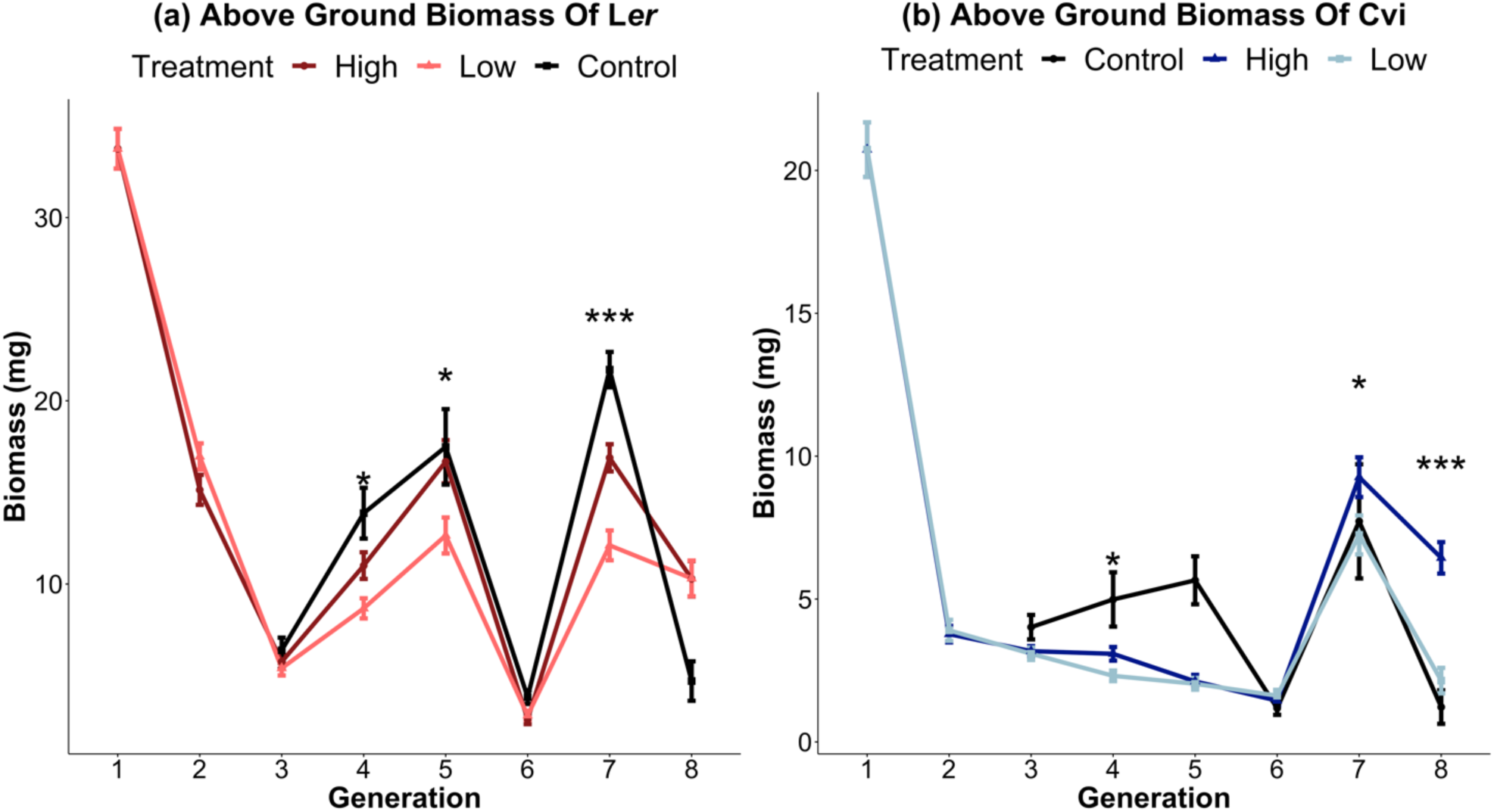
Ecosystem selection leads to changes in above-ground plant biomass. Above ground biomass for genotypes Ler (a) and Cvi (b) of *Arabidopsis thaliana*. Dark lines represent the high biomass treatment, and light lines represent the low biomass treatment. Each point represents the mean of n=50 microcosms. The mean biomass of the selection lines differed for several generations of both genotypes (t.test p-value * < 0.05; **<0.01;*** <0.001).

**Figure 2.**
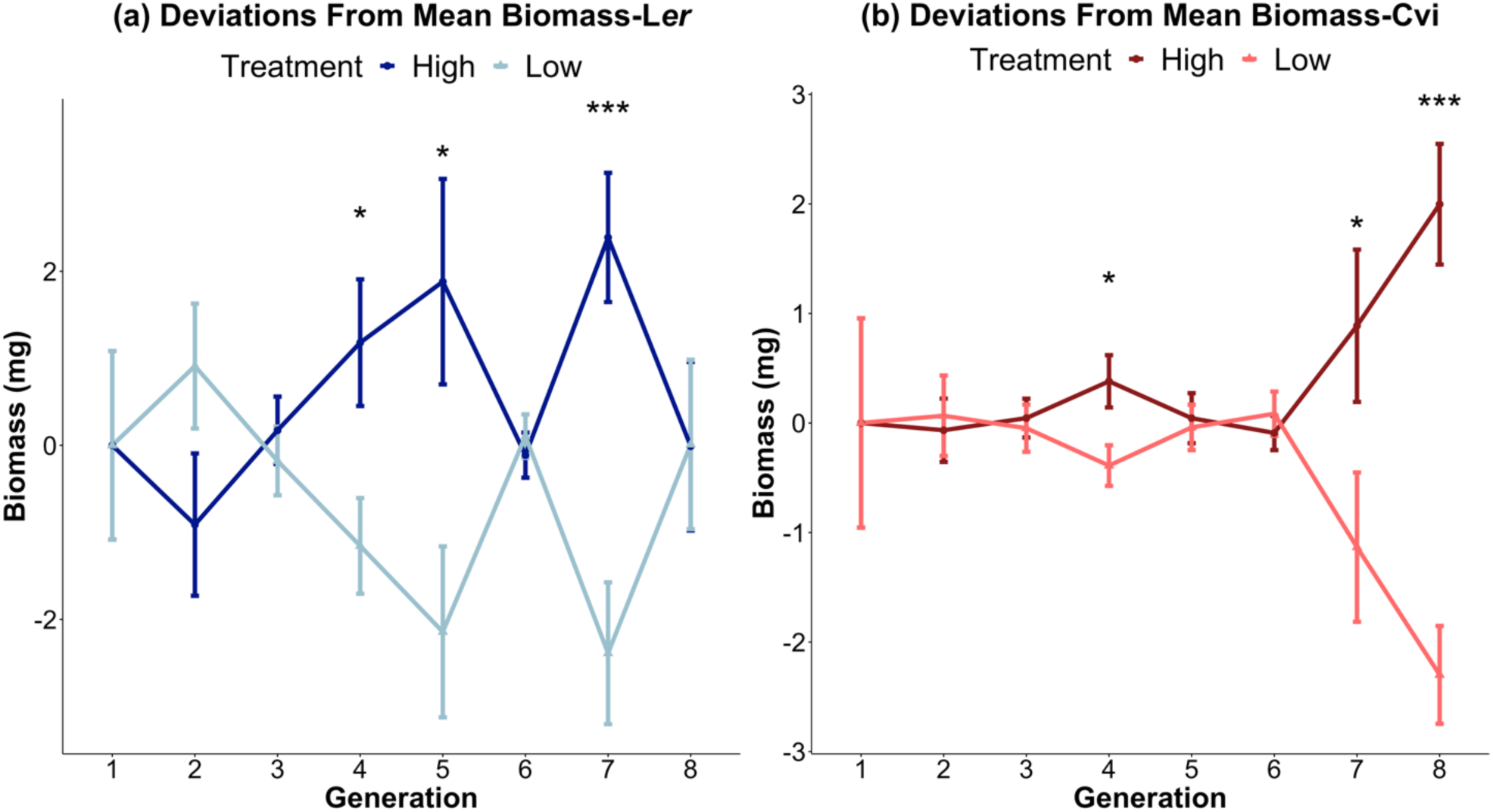
Above ground biomass represented as deviations from means for each generation. By plotting deviations from the mean, the influence of REVs on above-ground biomass is mitigated for genotypes L*er* (a) and Cvi (b). The dark lines denote high biomass treatments, while the light lines represent low biomass treatments. Each data point is the average of n=50 microcosms. (t.test p-value * < 0.05;**<0.01; *** <0.001).

### Microbial community Composition

After preprocessing the 16S rRNA metagenome amplicon reads in DADA 2, a total of 1,897,732 reads, with an average of 12822 reads per sample were obtained. Singletons and chimeras were removed during pre-processing. The most well-represented phyla were *Proteobacteria* (36.2%), *Bacteroidetes* (10.3%), *Planctomycetes* (7.8%), and *Actinobacteria* (7.6%) (Supplementary Figure S2). The most well-represented Classes identified in the dataset were *Alphaproteobacteria* (24.7%), *Gammaproteobacteria* (11.4%), *Bacteroidia (10.1%)*, and *Verrucomicrobiae* (6.2%) (Figure 3). The relative abundance is typical of microbiome data, heavily weighted to a few abundant groups with a long tail of rare taxa.

**Figure 3.**
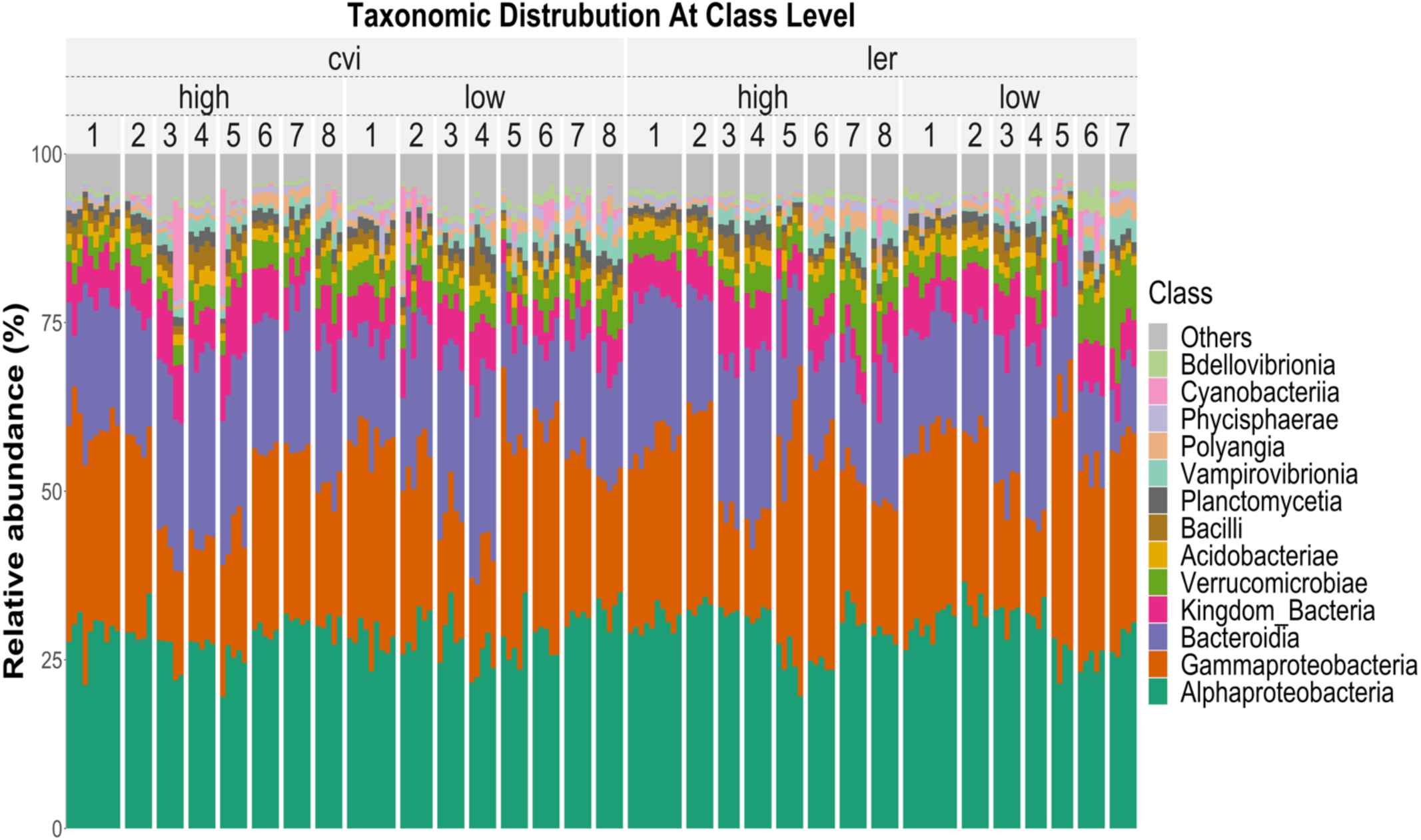
Taxonomic distribution of the microbial community is represented in terms of the relative abundance at the Class level. The plot shows eight generations (1-8) for the high and low biomass treatments of both the Cvi (left) and L*er* (right) genotypes of *Arabidopsis thaliana*. Ten samples were assayed for generation 1 experiments while five were done for all subsequent generations. The most represented classes across all samples were *Alphaproteobacteria* (24.7%), *Gammaproteobacteria* (11.4%), *Bacteroidia* (10.1%), and *Verrucomicrobiae* (6.2%).

### Alpha Diversity

We used two metrics to assess alpha diversity (Figure 4). Observed (F_7,160_ =333.054, P<0.001) estimates species richness, and showed a statistically significant decrease in the number of OTUs (1477 to ∼580) and richness across generations. The Shannon diversity index (F_7,160_ =82.23, P<0.001), which is more sensitive to the difference in abundance, also showed this steady decrease across generations. A negative exponential function was fitted to the Observed metric (R2=0.81, P<0.01) and the Shannon metric (R2=0.58, P<0.01). The Observed metric displayed a better fit, indicating increased stability of the microbiome in later generations. In addition, a linear mixed-effect model shows significant interactions between genotype:biomass treatment (F = 8.71, P < 0.01), genotype:generation (F =3, P < 0.01), and genotype:biomass treatment:generation (F = 3, P < 0.01), for Shannon and (Supplementary Table S5). No significant interaction terms were found for Observed, which does not consider the abundance of OTUs in its calculations.

**Figure 4.**
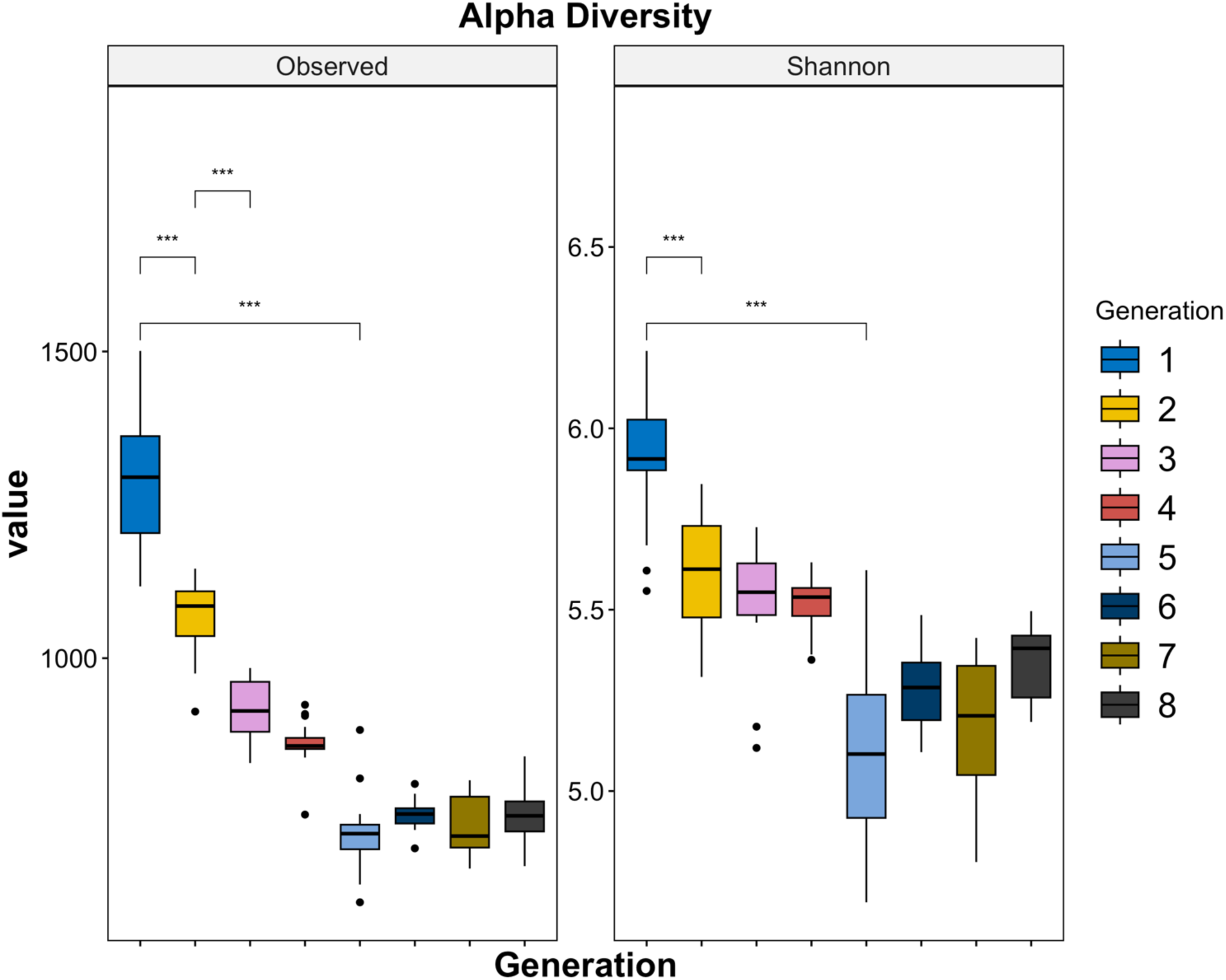
Changes in alpha diversity. Over the course of eight generations, two alpha diversity indices were computed, including the Observed and Shannon metrics. Results indicate a significant decline in the Observed number of OTUs, which estimates species richness. The Shannon index, which considers species abundance also exhibited a noteworthy decrease from generation 1 to 8. Additionally, a negative exponential function was fit to the Observed (R^2^=0.81, P<0.01) and Shannon (R^2^= 0.58, P<0.01) metrics, with the Observed showing a better fit indicating increased stability of the microbiome in later generations. (wilcox test *<0.05, **<0.01, *** <0.001).

### Beta Diversity

Principal coordinate analyses (PCoA) of the weighted UniFrac distances were plotted with the first two axes that captured the most variance in the data, from 73% in generation 1 to greater than 90% in generation 8 (Figure 5). UniFrac distance considers the evolutionary relationships between taxa, which makes it more biologically meaningful than other distance metrics that do not consider the evolutionary history. Weighted UniFrac is more sensitive to differences in the abundance of taxa. The microbiomes associated with the treatments and genotypes are very similar in generation 1, but begin to diverge by generation 2. By generation 3, the microbiomes of the L*er* and Cvi genotypes have diverged and by generation 5 the high and low biomass treatments appear to have further distinguished the microbial communities. The divergence of the microbial community in generations 4 and 5 caused by the high/low biomass treatments also aligns with the onset of notable differences in the above-ground plant biomass within these generations.

**Figure 5.**
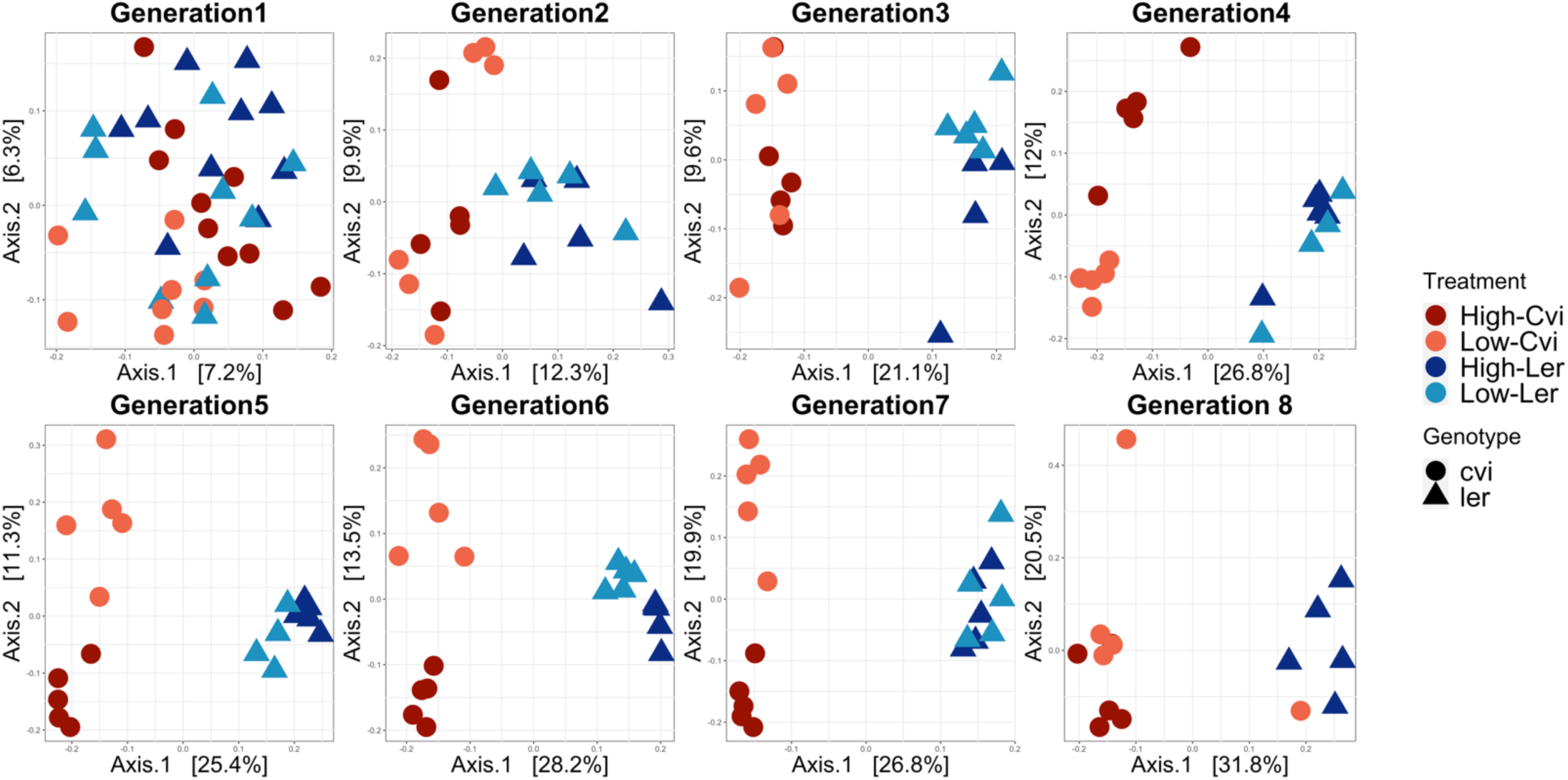
Host genotype and biomass treatment shape the composition of the microbiome. **Principle** inate Analysis (PCoA) using unweighted UniFrac for generations 1 through 8. Each point represents an dual sample with microbial communities defined of Cvi represented by red/orange circles and L*er* by ight blue triangles. The first two coordinate axes plotted accounted highest variation in the data, g from 13% in generation 1 to approximately 52% in generation 8. While there was no discernible ring in generation 1, both genotypes and high/low biomass treatments exhibited greater clustering over urse of the experiment.

The complex interplay between genotype and biomass treatments was modeled using a PERMANOVA test with both weighted and unweighted UniFrac distances to account for differences in abundance and presence-absence of OTUs respectively. The resulting R^2^ values for genotype and biomass treatment and residuals as a proxy for REVs were plotted (Figure 6a and 6b). For both weighted and unweighted UniFrac, the variance in microbial community explained by both genotype and biomass treatment increased significantly over the eight generations. Despite observing a decrease in the variance accounted for by the residuals, which could potentially signify a decline in the influence of REVs, they accounted for ∼50 percent of the variability within the microbial community in generation 8. For all metrics, weighted UniFrac exhibits more fluctuations compared to the unweighted UniFrac. Weighted UniFrac analyses found that genotype (F_1,39_=1.55, P<0.05) and biomass treatment (F_1,39_=1.33, P<0.001) explained 5.7% and 13.7%, respectively, of the dissimilarity between microbiomes in generation 1. By generation 8, genotype (F_1,14_=5.14, P<0.05) and biomass treatment (F_1,14_=5.48, P<0.001) explain 24% and 22%, respectively, of the dissimilarity between microbiomes (Supplementary Tables S6 and S7). For unweighted UniFrac, the influence of genotype on the microbial community increased from 3.3% in generation 1 (F_1,39_=1.33, P<0.01) to 26% in generation 8 (F_1,39_=5.13, P<0.001), and for biomass increased from 3.8% in generation 1 (F_1,14_=1.55, P<.001) to 13% in generation 8 (F_1,14_=2.58, P<0.05) (Supplementary Tables S8 and S9).

**Figure 6.**
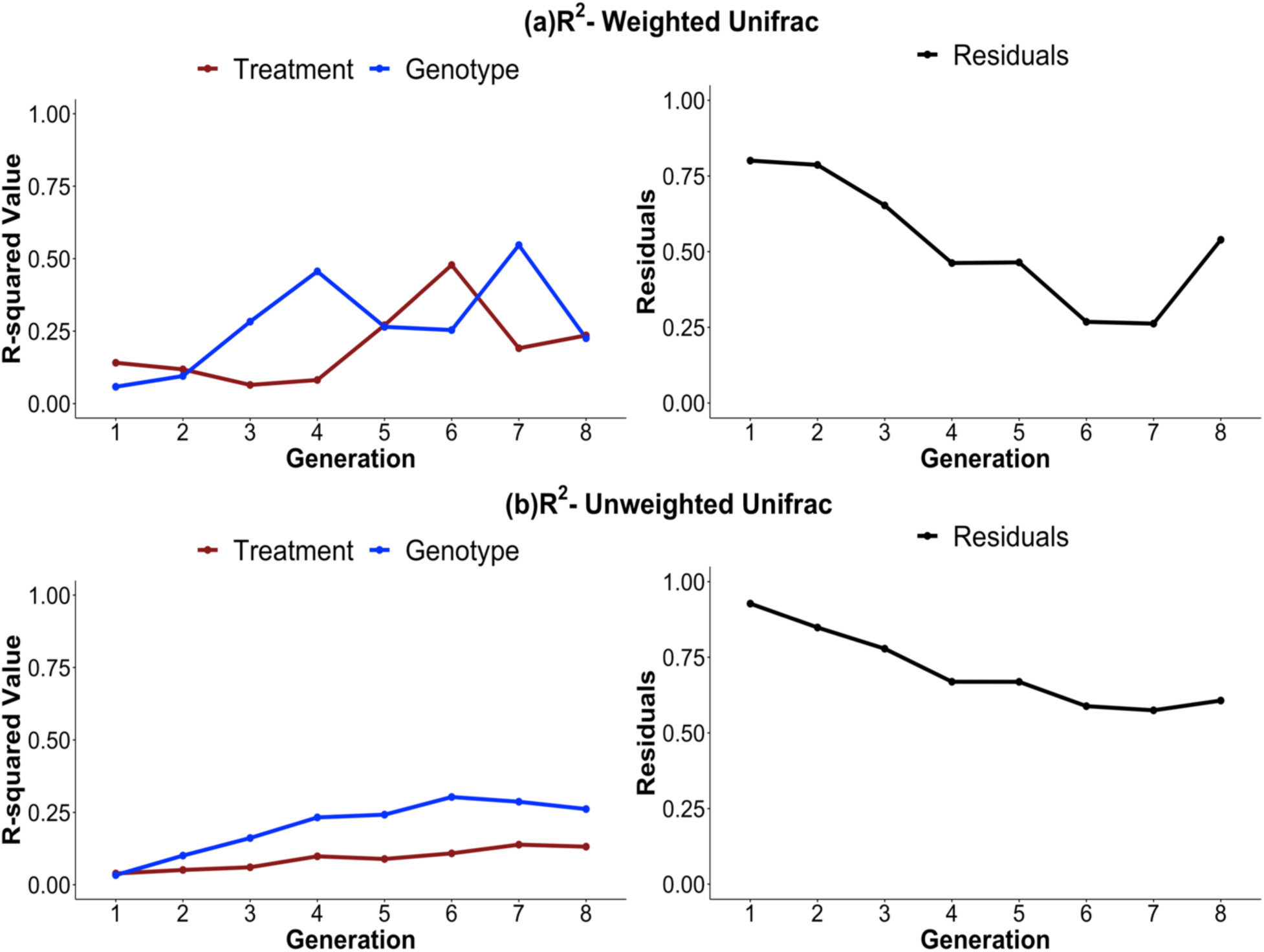
Variance in microbiome structure and composition explained by genotype and biomass treatment increases over generations. The R^2^ values were computed using the PERMANOVA test with the adonis2 function for generations 1 through 8 for (a) weighted and (b) unweighted-UniFrac distances. The resulting R^2^ and variance explained by residuals values for the biomass treatment (brown), genotype (blue) and residuals (black) are plotted. Residuals are plotted as a proxy for random environmental variations (REVs). There is a marked increase in the influence of both genotype and biomass treatment on differences in the microbial community from generations 1 through 8. Despite observing a decrease in the variance accounted for by the residuals, which could potentially signify a decline in the influence of REVs, the model elucidates more than 50 percent of the variability within the microbial community in the 8th generation. These differences are more pronounced in the weighted vs unweighted UniFrac. Model (distance.matrix∼biomass treatment +genotype).

### Sloan Neutral Model

Many forces can alter microbial community structure and dynamics and in our study these can be divided into selective forces such as the plant genotype being colonized and the biomass selection process, or neutral, stochastic forces innate in any experiment and environment. The Sloan neutral model was fit to the data to assess the importance of selective versus neutral drivers of change in the experiment. If neutral processes were the driving force in the microbial community assembly, the null hypothesis would be that all generations of the experiment would fit the model equally well. The model fit to neutrality represented by the R^2^ value decreases from 0.795 in generation 1 to 0.59 in generation 8 demonstrating the increasing importance of selective drivers through the course of the experiment (Figure 7).

**Figure 7.**
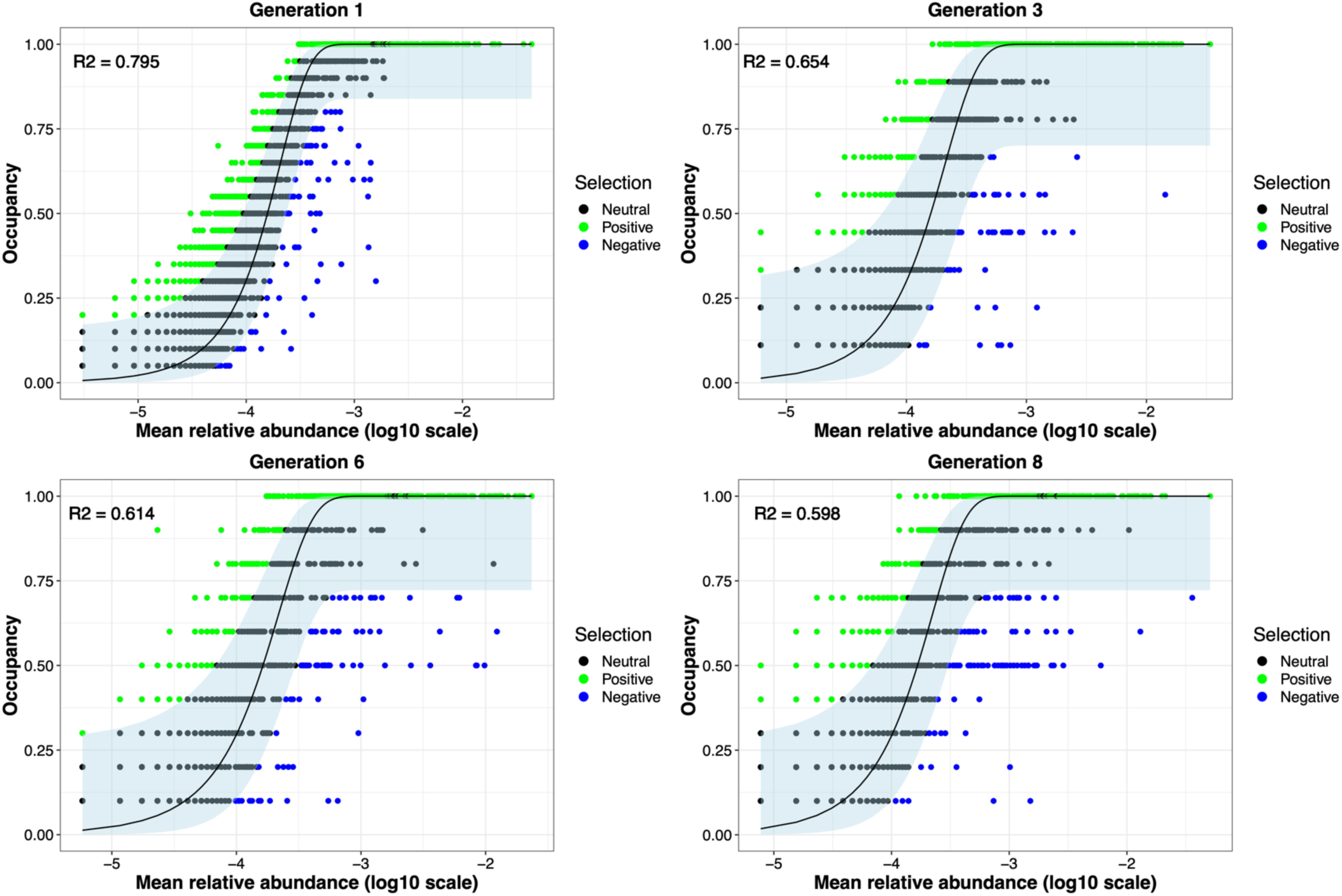
Fit of the Neutral model diminished as generations progressed. The occupancy (prevalence of OTU in) plotted against log(10) abundance is depicted using the Burns model. OTUs that are neutral, i.e., not for or against, are indicated in black. Green and blue-colored OTUs signify positive and negative selection, respectively. This plot highlights the presence of numerous microbes that undergo directed selection throughout the experiment.

### Differential Abundance

Differentially abundant (DA) taxa that distinguished between the high and low biomass treatments were determined using ANCOMBC2 at the Family level. Taxa lacking classification at the Family level were denoted by the subsequent identifiable taxonomic tier. The OTUs that were present in higher (red) and lower (blue) abundance in the high biomass treatment are identified (Figure 8a and 8b). Among the DA taxa in this study, several are known to benefit plants. The Class *Bacilli* promote plant growth [48]. The Class *Gemmatimonadetes* are positively associated with vegetation restoration, plant richness, and soil nutrients [49].

*Cytophagacea* are chemoorganotrophs and important for remineralizing organic materials into micronutrients. They could support both mycelial growth and plant nutrition [50].

**Figure 8.**
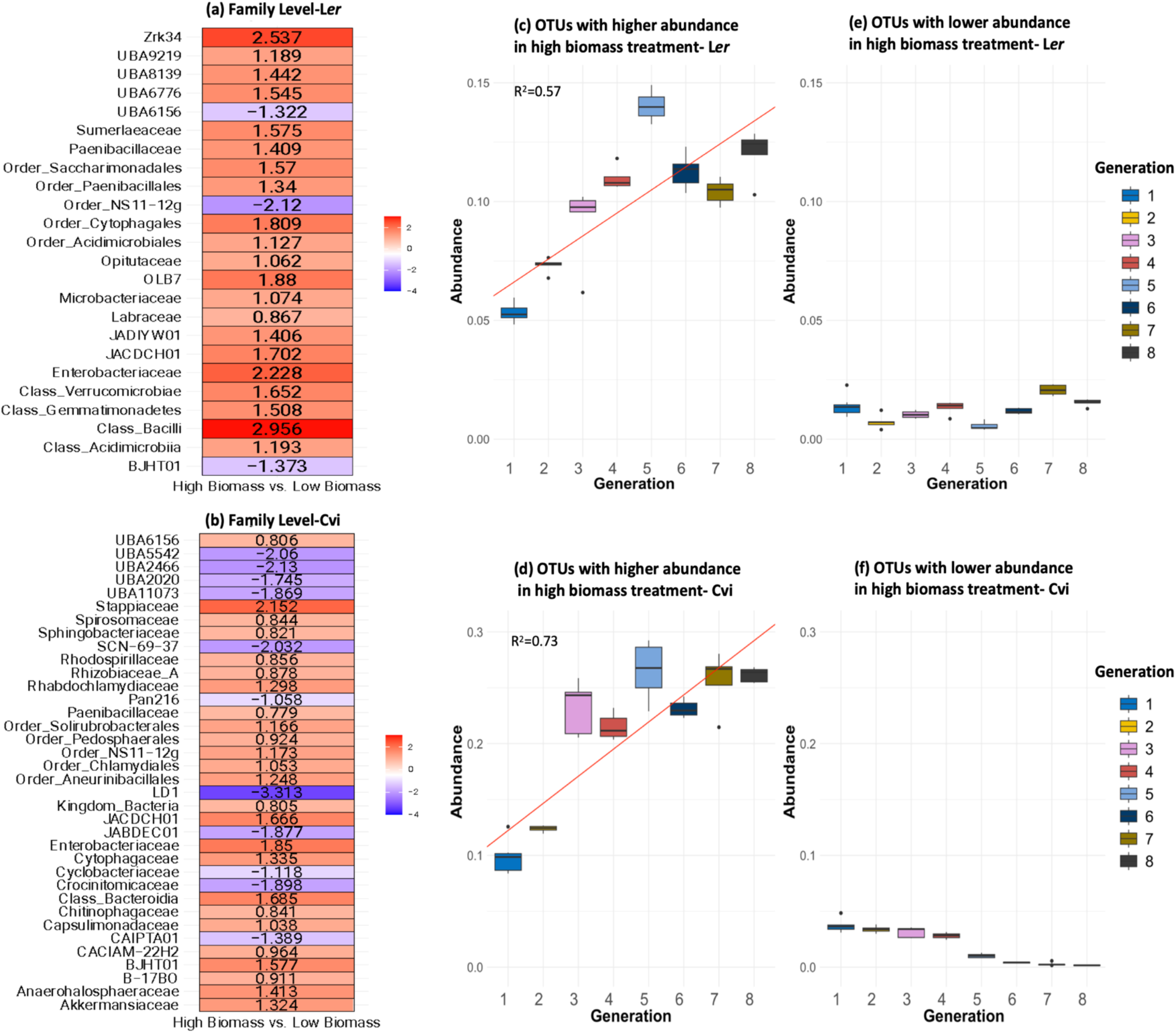
Differentially abundant OTUs. ANCOMBC2 results are illustrated as log_2_ fold changes at the family level for ss vs low biomass samples. Each taxon listed in the figure represents a single OTU in either the L*er* (a) or Cvi al community identified to the family level if possible. The scale indicates log_2_ fold change, red and blue for positive and negative fold change in the high biomass samples respectively. Well-known plant growth-promoting bacteria, such as *Paenibacillaceae, Bacilli, Labraceae, Rhizobiaceae* and *Bdellovibrio* are present in higher abundance. A vast majority of the other taxa identified are uncultivated and have little to no information on them. Figures (c) and (d) show box plots of relative abundance for all OTUs that were enriched in samples of the high biomass treatment in genotype L*er* and Cvi, respectively, with colors indicating different generations. These OTUs demonstrate a consistent trend of increasing relative abundance across successive generations. To test this trend (red line), a linear model was fitted to the data, resulting in an R-squared value of 0.57 (P < 0.01) for L*er* and 0.73 (P < 0.01) for Cvi. Figures (e) and (f) are box plots of relative abundance for all OTUs that were depleted in samples from the high biomass treatment in genotypes L*er* and Cvi, respectively, and colored by generation.

*Enterobacteriaceae*, *Paenibacillaceae*, and *JACDCH01* were all present in higher abundance in the high biomass treatments of both L*er* and Cvi. Most *Paenibacillaceae* members predominantly inhabited soil, frequently in close association with plant roots [51]. These rhizobacteria were known to play a significant role in enhancing plant growth and possessing potential applications in agriculture and *Enterobacteriaceae* had been reported to enhance plant growth [52–54].

Additionally, *Order_NS11-12g*, *BJHT01*, *JACDCH01*, and *UBA6156* were more abundant in Cvi under high biomass conditions and less abundant in L*er* under high biomass conditions. The majority of these were uncultivated or candidate taxa. Two alternative families in the Order *Rhizobiales* appeared differentially associated with the two plant genotypes: *Labraceae* with L*er* and *Rhizobiaceae* with Cvi [55]. These could indicate genotype-specific interactions with different members of the microbial community.

The relative abundance of all OTUs that exhibited higher abundance (enriched) in samples from the high biomass treatment in the L*er* and Cvi is depicted in Figures 8 (c) and 8 (d) respectively. These OTUs consistently exhibit an increasing trend in relative abundance across successive generations. A linear model was employed to test this trend, resulting in R-squared values of 0.57 (P < 0.01) for L*er* and 0.73 (P < 0.01) for Cvi. Furthermore, OTUs that were lower in abundance (depleted) in samples from the high biomass treatment in L*er* and Cvi are depicted in Figures 8 (e) and 8 (f) respectively. These OTUs persist in low abundance across all generations, suggesting a lack of selective pressure on this group of microbes in the samples from the high biomass treatment.

## Discussion

We applied artificial ecosystem selection for eight generations, in *Arabidopsis thaliana* L*er* and Cvi to select soil microbiomes associated with higher or lower biomass of the host. In contrast to some previous studies [27,33], we did not fertilize plants thus maintaining nutrient limitation, and thereby promoting the plant’s interaction with the microbiome [56]. Over the course of eight generations, a response to the selection, most apparent after generations 4 and 5, was observed both in the microbiome characteristics that shifted gradually and in the plant biomass. Microbiome selection noticeably influenced plant biomass despite large phenotypic variation from one generation to the next. Stochasticity in growth through generations may be attributed to random environmental variations (REVs) as indicated by Swenson et al. [28]. This was observed in the uninoculated sterile potting soil reference that showed similar patterns of variability as the inoculated treatments across generations (Figure 1). Ecological variability among generations even under controlled growth chamber conditions is a common characteristic of studies [57] and may be due to minute fluctuations in growth chamber conditions or batch and age of potting soil.

We observed a gradual decline in microbial species richness during the initial generations of the experiment. This was evident from the observed OTU counts, as well as the Chao1 and Shannon diversity indices. This pattern is consistent with findings from other studies [32–34], highlighting the impact of selection pressures as the microbial communities adapted to the host plant environment. However, in our study which continued for about twice the number of generations of these earlier studies, we observed a stabilization of richness and alpha diversity in the later generations. This was further substantiated by the stronger fit to the negative exponential function when compared to a linear model, indicating constant and proportional changes in alpha diversity and the rhizosphere microbiome’s trend towards reduced diversity loss over time, accompanied by heightened stability over eight generations.

To further understand the complex interplay between the forces of directed selection (plant genotype and biomass treatment), we did a PCoA. Over successive generations, the results demonstrate a strengthening effect of plant genotypes and biomass treatment on the microbial community (Figure 5). This result was modeled using PERMANOVA, for both weighted and unweighted UniFrac distances. Both weighted and unweighted UniFrac show marked increases in the proportion of variance explained by genotype and biomass treatment, with unweighted UniFrac exhibiting larger increases from generation to generation (Figure 6). Both metrics consider phylogenetic relatedness. Weighted UniFrac considers the absolute abundance of OTUs and is generally used to study changes in microbial community structure. Unweighted UniFrac considers only the presence/absence of OTUs and is generally used to study changes in microbial community composition. The results suggest that variance in microbiome composition as explained by genotype consistently increases, whereas variance in microbiome structure as explained by genotype, fluctuates more with changes in abundant taxa.

Here we would like to highlight that pronounced shifts in biomass between the high/low biomass treatments, alpha diversity, and beta diversity all occur around generations 4 and 5 (Figure 1 and 2). This marks the point at which the changes in alpha diversity stabilize (Figure 4) and the divergence between microbial communities by high/low biomass treatments in the PCoA plots becomes more pronounced (Figure 5). It is possible that over the first four to five generations, the starting microbial community underwent a period of restructuring before it stabilized and formed four distinct communities under selection by high/low biomass treatments and by plant genotype. We propose a complex interaction between plants and their associated microbiome ensued, where differences in root exudation patterns between the two genotypes presumably established associations with microbes in the early generations. Simultaneously, during this period, the biomass treatment likely promoted host-specific microbe-mediated interactions that modulate plant biomass. This restructuring of the microbial community is driven by both structural (abundance of OTUs) and compositional (presence/absence of OTUs) effects in the microbiome (Figure 6a and 6b). Differences in the abundance of taxa driven by the biomass and genotype selection pressures, was a larger contributor to the generation-to-generation variation during the experiment.

Changes in the assembly of microbial communities can arise from either selective pressure, like the plant genotype or biomass treatment observed in this experiment, or stochastic processes, like minute changes in growth chamber humidity or microenvironment. To gain deeper insights into the influence of selection on microbial community assembly, a neutral model was fit to the data [44,45]. Interestingly, this revealed a progressive decline in fit to neutrality, indicated by decreasing R^2^ values and increasing AIC values (Figure 7). This suggests an increasing influence of selective forces such as biomass treatment and genotype over multiple generations of ecosystem selection. This was also previously observed by Morella et al. [34].

Previous studies in *Arabidopsis thaliana* have often detected only a weak genotypic effect on the rhizosphere microbiome [23,24,58]. However, a majority of these studies are focused on a single growth cycle. Insights from plant-soil feedback research emphasize that the interactions between plants and their microbiomes can be modified by consecutive growth cycles of the same plant species in its respective soil [25,26]. Recent research has presented conflicting findings on the influence of host genotype [33,34,59]. The different experimental designs, communities being assessed and the length of these experiments, alongside high stochastic environmental variation that is clearly associated with these microbial studies likely account for the conflicting findings. In our investigation, we find that during the initial stages, genotype and biomass treatments have modest but significant impacts. Over time, the plant genotype and biomass treatments have an increasing influence. Together these selective forces explained ∼40% of the variation in the microbial community composition in these later generations.

A key aim of microbiome engineering has been to develop host-specific microbial communities that impart lasting beneficial effects on the plant host [27,29,32,60]. Here we show the enrichment of some common taxa in high biomass treatments but also some genotype-specific changes. Despite starting with the same soil, only three common families were enriched in the high biomass treatment samples of both the L*er* and Cvi genotypes. In addition, well-known plant growth-promoting rhizobacteria were enriched in a genotype-specific manner for both L*er* and Cvi high biomass treatment samples. In the Order *Rhizobiales*, *Labraceae* was enriched in L*er,* while *Rhizobiaceae* was enriched in Cvi [55]. Moreover, in high biomass conditions, Cvi had increased levels of the taxa Order_NS11-12g, BJHT01, JACDCH01, and UBA6156, while L*er* had decreased levels of these. These data suggest genotype-specific plant genetic control of the rhizosphere microbiome.

To further explore how the abundance of these OTUs changed across successive generations, we grouped all the DA OTUs that were enriched or depleted in the samples of high biomass treatment and calculated the sum of their relative abundance. The OTUs enriched in the samples of high biomass treatment consistently displayed an increasing trend in relative abundance over subsequent generations (Figures 8c and 8d). A linear model was fitted to this data with high R^2^ values, reinforcing the notion that the genotype and biomass treatments exerted selective pressure on entire microbial groups, rather than individual species. Conversely, the OTUs that were depleted in the samples of high biomass treatment consistently remained in a state of low relative abundance throughout the entirety of the experiment (Figures 8e and 8f).

This observation may indicate that the selective pressures exerted by the genotype and biomass treatments do not influence this particular group of microbes.

Taken collectively, these findings suggest the presence of diverse selection pressures influences the rhizosphere microbiome. Initially, it seems that neutral processes seem to play a major role in determining the structure, and composition of the microbiome (Figure 7). As generations progress, we observe an increase in the strength of directional selection, possibly driven by genotype and biomass treatment, resulting in the enrichment of specific groups of microbes, including some PGPRs (Figures 8c and 8d). The persistence of depleted microbes in high biomass treatment samples suggests the possibility for negative frequency-dependent selection, preserving them at a low abundance. On the other hand, it is possible that this group is experiencing no observable selection pressure.

Multiple studies have shown variations in the structure of the rhizosphere microbiome, even within closely related plant genotypes, highlighting the importance of genotype-specific root exudates in forming associations with the corresponding microbiome [61–65]. Further exploration must prioritize highlighting the mechanisms that uphold these interactions through the utilization of comparative metagenome, metatranscriptome and metaproteome analyses. This will play a part in decoding chemical crosstalk, which could eventually foster interactions between plants and microbes, leading to an overall enhancement in plant fitness.

The relationship between plants and microorganisms in the rhizosphere involves complex and diverse interactions, which influence crucial ecological and physiological processes. These interactions can be mutualistic, competitive, or antagonistic in nature. Many biotic and abiotic factors act in concert to influence the dynamic rhizosphere microbiome. Gaining insights into how these factors influence the assembly of microbial communities is crucial for obtaining targeted and long-lasting advantages for plants. Nevertheless, establishing a strong and host-specific microbial community that consistently provides beneficial effects continues to be a persistent challenge. We have shown that, despite stochastic fluctuation due to REVs, it is possible to select for microbial communities that impact biomass in a host-specific manner within four generations, addressing this key challenge in microbial community engineering [32,60]. The rhizosphere microbiome that evolves under plant-mediated selection offers better chances of survivability and efficacy when applied as inoculum to the plant [66,67]. Multi-species ecosystem selection provides a perspective on microbiome assembly that indicates the holobiont, and the evolving relationship between host and microbiome contributes to emerging properties beyond those predicted by host genotype or initial microbial community composition. This study enhances our understanding of the temporal dynamics involved in microbial assembly in the rhizosphere and has implications for sustainable agriculture, evolution, and ecology.

## Supplementary Materials

Figure S1: Rhizotubes schematic; Figure S2: Taxonomic distribution of the microbial community represented in terms of the relative abundance at the Phylum level. Table S1: Statistical comparisons of biomass for L*er*; Table S2: Statistical comparisons of biomass for Cvi; Table S3: ANOVA for Observed alpha diversity index; Table S4: ANOVA for Shannon alpha diversity index; Table S5: Linear mixed effect model for Shannon diversity index; Table S6: PERMANOVA weighted UniFrac distances for generation 1; Supplementary Table S7: PERMANOVA weighted UniFrac distances for generation 8; Supplementary Table S8: PERMANOVA unweighted UniFrac distances for generation 1; Table S9. PERMANOVA unweighted UniFrac distances for generation 8.

## Author Contributions

N.S. Conceptualization, methodology, data analysis and visualization, data collection, writing-original draft, writing-reviewing and editing. P.S. Methodology, data analysis, writing-original draft, writing-reviewing, and editing. T.M. Data collection. R.K. Resources, funding, writing-reviewing, and editing.

## Data Availability Statement

All sequence data and metadata can be downloaded from PRJNA1010286.

## Acknowledgments

We would like to express our sincere gratitude to the following individuals for their advice at various stages of this project Professor David Sloan Wilson, Dr. Jenny Kao-Kniffin, Dr. Deepa Agashe, Dr. Gergely Maróti, Professor Douglas Woodhams, Professor Emeritus Michael Shiaris, Dr. Karen Ricciardi, Dr. Patrick Kearns, Dr. Kourosh Zarringhalam, Dr. David Degras-Valabregue, Professor Liam Revell and Professor Jay T. Lennon, Dr. Molly Bletz, Dr. Brandon LaBumbard, and Cassandra Lyons.

