## supplementary for "Multi-Generation Ecosystem Selection Of Rhizosphere Microbial Communities Associated With Plant Genotype and Biomass In *Arabidopsis thaliana*"

### SUPPLEMENTARY FIGURES AND TABLES

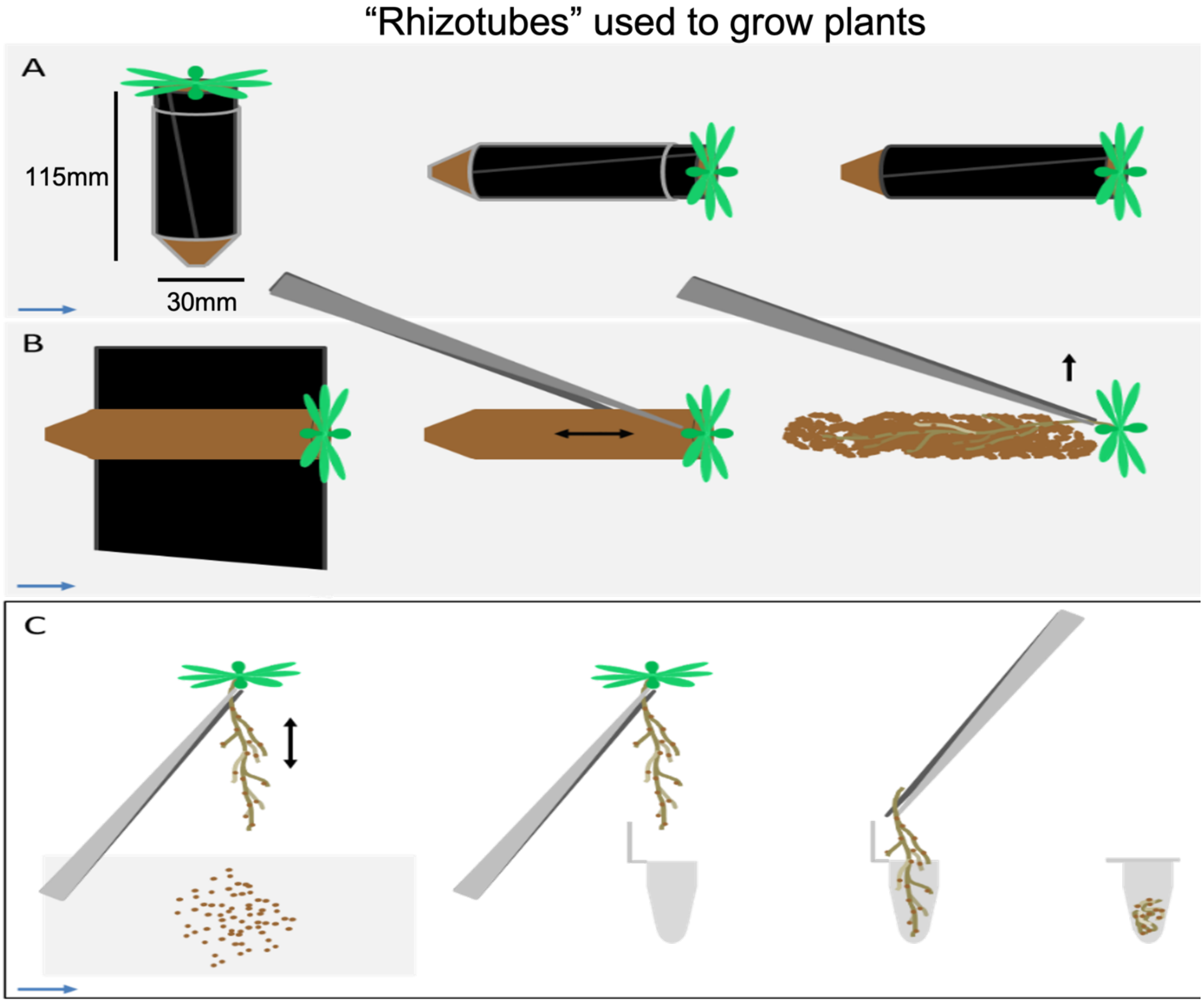


**Supplementary Figure S1. Rhizotube shchematic.** The purpose of this illustration is to demonstrate how the custom-made rhizotubes facilitate the collection of plant rhizosphere. The black insert within the tube is unwrapped to extract the rhizosphere, which is then separated from the bulk soil by shaking the roots. After removing the above-ground portion of the plant, the root-soil complex (comprising the rhizosphere and the endosphere) was obtained by shaking off the excess soil and then placed in a sterile 5ml tube and bulk soil samples were moved to a sterile Ziplock bag. Both were immediately transferred to dry ice and then stored at -80oC for DNA analysis.

.

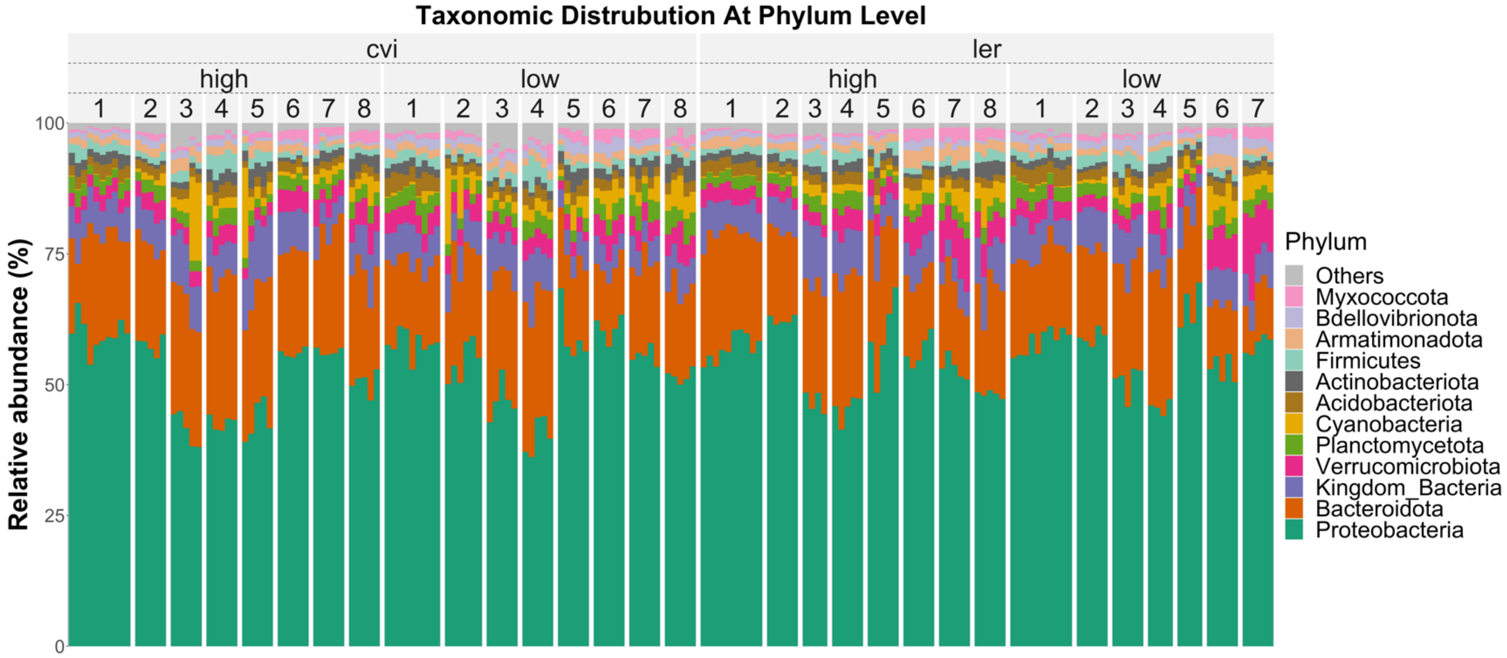


**Supplementary Figure S2. Taxonomic distribution of the microbial community represented in terms of the relative abundance at the Phylum level.** The plot shows eight generations (1-8) for the high and low biomass treatments of both the Cvi (left) and L*er* (right) genotypes of *Arabidopsis thaliana*. The microbial community is dominated by *Proteobacteria*, *Bacteroidetes*, *Verrucomicrobia, and Planctomycetes* which together comprise nearly 73% of the bacterial community.

| Generation | group1 | group2 | p | p.signif |
|---|---|---|---|---|
| Gen4 | High | Low | 1.222020e-02 | * |
| Gen5 | High | Low | 1.032381e-02 | * |
| Gen7 | High | Low | 3.562072e-05 | **** |

**Supplementary Table S1 Statistical comparisons of biomass for L*er*** samples (t.test p-value * < 0.05; **<0.01; *** <0.001).

| Generation | group1 | group2 | p | p.signif |
|---|---|---|---|---|
| Gen4 | High | Low | 1.275260e-02 | * |
| Gen7 | High | Low | 4.123576e-02 | * |
| Gen8 | High | Low | 6.758467e-08 | **** |

**Supplementary Table S2 Statistical comparisons of biomass for Cvi** samples (t.test p-value * < 0.05; **<0.01; *** <0.001).

```
              Df Sum Sq Mean Sq F value Pr(>F)    
Generation     7 158.13  22.590 333.054 <2e-16 ***
Lineage        1   0.01   0.009   0.130  0.719    
Regime         1   0.01   0.011   0.169  0.681    
Residuals    160  10.85   0.068                   
---
Signif. codes:  0 ‘***’ 0.001 ‘**’ 0.01 ‘*’ 0.05 ‘.’ 0.1 ‘ ’
```

**Supplementary Table S3. ANOVA for Observed alpha diversity index** shows a significant influence of generation on alpha diversity. Lineage is genotype and Regime is biomass treatment. Design (Observed ~ Generation + Lineage + Regime) Df: degrees of freedom; SumOfSqs : sum of squares; Mean sq: mean squares; F: F statistic; Pr(>F): p-value.

```
              Df Sum Sq Mean Sq F value Pr(>F)    
Generation     7 132.11  18.873  82.238 <2e-16 ***
Lineage        1   0.01   0.009   0.040  0.841    
Regime         1   0.16   0.160   0.695  0.406    
Residuals    160  36.72   0.229                   
---
Signif. codes:  0 ‘***’ 0.001 ‘**’ 0.01 ‘*’ 0.05 ‘.’ 0.1 ‘ ’
```

**Supplementary Table S4. ANOVA for Shannon alpha diversity index** shows a significant influence of generation on alpha diversity. Lineage is genotype and Regime is biomass treatment. Design (Observed ~ Generation + Lineage + Regime) Df: degrees of freedom; SumOfSqs : sum of squares; Mean sq: mean squares; F: F statistic; Pr(>F): p-value.

Type III Analysis of Variance Table with Satterthwaite's method

| | Sum Sq | Mean Sq | NumDF | DenDF | F value | Pr(>F) | |
|---|---|---|---|---|---|---|---|
| Lineage | 0.0035 | 0.0035 | 1 | 139 | 0.0065 | 0.936050 | |
| Regime | 0.0945 | 0.0945 | 1 | 139 | 0.1770 | 0.674654 | |
| Generation | 1.4097 | 0.2014 | 7 | 139 | 0.3771 | 0.914458 | |
| Lineage:Regime | 4.6552 | 4.6552 | 1 | 139 | 8.7162 | 0.003704 | ** |
| Lineage:Generation | 11.2206 | 1.6029 | 7 | 139 | 3.0013 | 0.005765 | ** |
| Regime:Generation | 2.8505 | 0.4072 | 7 | 139 | 0.7624 | 0.619688 | |
| Lineage:Regime:Generation | 9.7397 | 1.6233 | 6 | 139 | 3.0393 | 0.007990 | ** |

---

Signif. codes: 0 '***' 0.001 '**' 0.01 '*' 0.05 '.' 0.1 ' ' 1

**Supplementary Table S5. Linear mixed effect model for Shannon diversity index.** Shows a significant effect of the generation and interaction term with genotype ("Lineage") and biomass treatment ("Regime"). Design lmer (DiveristyMetric, Genotype*Generation*Regime, (1|Generation)) Sum Sq: sum of squares; Mean sq : mean squares; NumDF: numerator degrees of freedom; DenDF: denominator degrees of freedom; F value: F statistic; Pr(>F): p-value.

adonis2(formula = wunifrac ~ Regime + Lineage, data = meta, permutations = 999

| | Df | SumOfSqs | R2 | F | Pr(>F) | |
|---|---|---|---|---|---|---|
| Regime | 1 | 0.0020294 | 0.13730 | 6.3126 | 0.001 | *** |
| Lineage | 1 | 0.0008563 | 0.05794 | 2.6637 | 0.038 | * |
| Residual | 37 | 0.0118951 | 0.80476 | | | |
| Total | 39 | 0.0147809 | 1.00000 | | | |

---

Signif. codes: 0 '***' 0.001 '**' 0.01 '*' 0.05 '.' 0.1 ' ' 1

**Supplementary Table S6. PERMANOVA weighted UniFrac distances for generation 1.** The $R^2$ values were computed using the PERMANOVA test with the adonis2 function for generation 1 for weighted UniFrac distances. Genotype is referred to as "Lineage" and biomass treatment as "Regime". Model (distance.matrix~biomass treatment +genotype).

```
adonis2(formula = wunifrac ~ Regime + Lineage, data = meta, permutations = 999
         Df  SumOfSqs      R2      F Pr(>F)
Regime    1 0.0033556 0.22498 5.0755  0.006 **
Lineage   1 0.0036257 0.24309 5.4840  0.001 ***
Residual 12 0.0079337 0.53193
Total    14 0.0149149 1.00000
---
Signif. codes:  0 ‘***’ 0.001 ‘**’ 0.01 ‘*’ 0.05 ‘.’ 0.1 ‘ ’ 1
```

**Supplementary Table S7. PERMANOVA weighted UniFrac distances for generation 8.** The $R^2$ values were computed using the PERMANOVA test with the adonis2 function for generation 8 for weighted UniFrac distances. Genotype is referred to as “Lineage” and biomass treatment as “Regime”. Model (distance.matrix~biomass treatment +genotype).

```
adonis2(formula = unifrac ~ Regime + Lineage, data = meta, permutations = 999)
         Df SumOfSqs      R2      F Pr(>F)
Regime    1   0.2022 0.03895 1.5537  0.001 ***
Lineage   1   0.1740 0.03351 1.3369  0.006 **
Residual 37   4.8150 0.92754
Total    39   5.1912 1.00000
---
Signif. codes:  0 ‘***’ 0.001 ‘**’ 0.01 ‘*’ 0.05 ‘.’ 0.1 ‘ ’ 1
```

**Supplementary Table S8. PERMANOVA unweighted UniFrac distances for generation 1.** The $R^2$ values were computed using the PERMANOVA test with the adonis2 function for generation 1 for unweighted UniFrac distances. Genotype is referred to as “Lineage” and biomass treatment as “Regime”. Model (distance.matrix~biomass treatment +genotype).

```
adonis2(formula = unifrac ~ Regime + Lineage, data = meta, permutations = 999)
         Df SumOfSqs      R2      F Pr(>F)
Regime    1  0.20168 0.13113 2.5871  0.013 *
Lineage   1  0.40088 0.26065 5.1425  0.001 ***
Residual 12  0.93544 0.60822
Total    14  1.53799 1.00000
---
Signif. codes:  0 ‘***’ 0.001 ‘**’ 0.01 ‘*’ 0.05 ‘.’ 0.1 ‘ ’ 1
```

**Supplementary Table S9. PERMANOVA unweighted UniFrac distances for generation 8.** The $R^2$ values were computed using the PERMANOVA test with the adonis2 function for generation 8 for unweighted UniFrac distances. Genotype is referred to as “Lineage” and biomass treatment as “Regime”. Model (distance.matrix~biomass treatment +genotype).
